# Paranormal believers are quicker but less accurate in rejecting the presence of the target in conjunction visual search compared to skeptics

**DOI:** 10.1101/2024.04.25.590450

**Authors:** Fatemeh Akbari, Samaneh Asivandzadehchaharmahali, Abdolvahed Narmashiri

## Abstract

Recent studies have shown that paranormal believers may exhibit cognitive dysfunctions, yet their performance in conjunction with visual search has not been understood. To address this issue, we examined the performance of both paranormal believers and skeptics in a conjunction visual search task, paying particular attention to their search time and accuracy across different set sizes in both target-present (TP) and target-absent (TA) trials. In our study, believers demonstrated a tendency toward speed but also displayed carelessness compared to skeptics when rejecting the presence of the target. Conversely, skeptics exhibited slower search times but demonstrated greater accuracy both in rejecting the presence of the target and in finding it. Overall, our findings suggest that believers were quicker and less accurate in rejecting the presence of the target in conjunction visual search compared to skeptics, highlighting potential differences in cognitive processing between skeptics and believers.

**Significant statement:** Our study investigates the performance of paranormal believers and skeptics in conjunction with visual search tasks, shedding light on potential differences in cognitive processing between the two groups. While believers demonstrate faster search times, they also display greater carelessness compared to skeptics when rejecting the target’s presence. In contrast, skeptics exhibit slower search times but higher accuracy in both rejecting and finding the target. These findings underscore the importance of considering individual belief systems in understanding cognitive performance in conjunction with visual tasks.

## Introduction

Paranormal beliefs constitute a diverse spectrum of convictions, phenomena, and practices that challenge the foundational principles of scientific understanding ^1^. Within this broad framework, these beliefs traverse multiple domains, encompassing adherence to traditional religions alongside notions such as telekinesis, superstition, witchcraft, spiritualism, magical thinking, extrasensory perception, and precognition ^2^. This expansive range underscores the complexity and diversity of human beliefs and experiences that extend beyond conventional scientific explanations.

Research has explored the correlation between cognitive functioning and beliefs in the paranormal ^3-12^. Irwin ^13^ suggests that paranormal believers might experience cognitive deficiencies, a notion supported by several studies indicating shortcomings across various cognitive domains among this group. These deficits cover a range of cognitive functions, including inhibitory control ^3,14,15^, critical thinking ^16^, probability judgment ^17^, working memory, and inattention blindness ^18^, reasoning ^19^, and imagination ^20^. In addition, the role of an individual’s belief system in cognitive function cannot be overlooked. While religious and spiritual beliefs have been linked to slower cognitive decline in older adults ^21^, they have also been associated with lower memory performance and intelligence ^22^. Similarly, paranormal beliefs and conspiracy theories are connected to reduced analytical thinking and lower educational attainment ^23,24^. Such beliefs are also linked with tendencies towards illusory pattern perception ^4,5,25,26^, decreased cognitive reflection ^27,28^, biases against evidence ^29^, and hindsight bias ^30^. Despite these observed relationships, there remains uncertainty regarding how paranormal believers search to find a target and reject the presence of a target in conjunction with visual search tasks.

To address this issue, we conducted a behavioral experiment using a conjunction visual search. This task comprised two types of trials: those with a target presented in the array (target-present trials) and those without a target in the array (target-absent trials). We examined whether paranormal believers exhibit differences in search time and accuracy in recognizing significant target findings and in rejecting the presence of the target amidst set size levels ranging from 3 to 9 in the conjunction visual search. Furthermore, believers demonstrated a tendency toward faster search times but also displayed carelessness compared to skeptics when rejecting the presence of the target. Conversely, skeptics exhibited slower search times but showcased greater accuracy both in rejecting the presence of the target and in finding it in conjunction visual search.

## Method

### Participants

The experiment involved 46 participants (6 males, 40 females; mean age: 39.55 years). The severity of paranormal beliefs was determined using the overall score on the Paranormal Belief Scale (PBS) 36. On the PBS, the overall average score was 36.95 (SD = 7.46). After conducting a median split (median = 38, see Fig 1B), two groups were formed: paranormal believers (n = 23; 3 males, 20 females; mean age = 38.81 years; mean PBS score = 42.82, SD = 2.80) and skeptics (n = 23; 3 males, 20 females; mean age = 40.21 years; mean PBS score = 31.08, SD = 5.83). All participants reported normal or corrected-to-normal vision and underwent screening for mental illnesses, personality issues, drug or alcohol abuse, neurological disorders, and epilepsy. Participants were compensated with gifts for their involvement.

**Fig 1.**
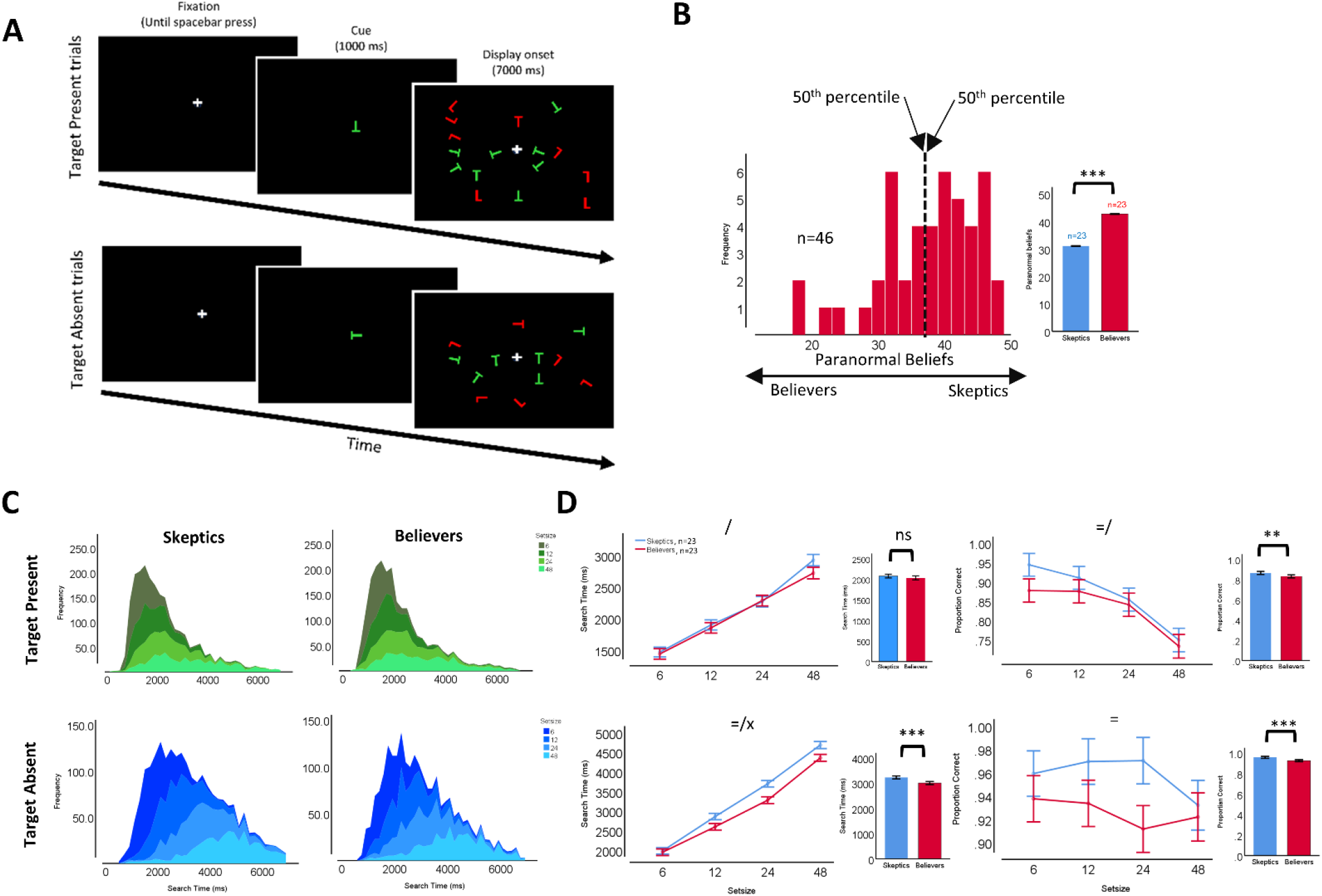
Visual Search Paradigm, Task Design, and Behavioral Results. A) The search task comprised Target-Present (TP) and Target-Absent (TA) trials. In each trial, a variable number of objects (6, 12, 24, or 48) were presented on the screen. In TP trials, only one of the displayed objects served as the target, while in TA trials, there was no target. Subjects were required to press two buttons to indicate the presence or absence of the target within the array. The task required an initial fixation (until the spacebar press) on a fixation at the screen center followed by a brief presentation of the target for that trial (1000 ms), followed by a presentation of the search array (7000 ms). B) Paranormal beliefs score distribution for all participants. The x-axis shows the paranormal belief scores, and the y-axis indicates the frequency. Paranormal belief scores in the lower 50th percentile or higher 50th percentile were grouped as believers and skeptics, respectively. Insets bar plots show the average paranormal belief scores for believers (red) and skeptics (blue) based on the percentile definition. C) Search time distribution for TP trials (top, depicted in a green gradient color) and TA trials (bottom, represented in a blue gradient color) across various set sizes for believers (right) and skeptics (left). D) Search time (in ms, on the left) and accuracy (in %, on the right) as functions of set size for TP trials (depicted in purple) and TA trials (represented in blue). E) Search time (in ms, at the left) and accuracy (in %, at the right) for TP trials (top) and TA trials (bottom) across collapsed set sizes for believers (red) and skeptics (blue). Insets bar plots show the average search time (left) and accuracy (right) for TP trials (top) and TA trials (bottom) for believers (red) and skeptics (blue). The symbols *, **, and *** indicate statistical significance levels at p < 0.05, p < 0.01, and p < 0.001, respectively. The symbols =, /, and x represent the main effects of search efficiency, set size, and their interaction, respectively.

## Materials

### Paranormal Belief Scale (PBS) ^31^

This paranormal belief scale includes 10 statements about paranormal beliefs, such as “I have had at least one experience of having a relationship between my thoughts and those of another person.” Participants were asked to choose the option that best fit them. Lower scores indicate stronger beliefs in paranormal phenomena. The scale consists of 10 questions with 5 options ranging from “completely agree” to “completely disagree.” Lower scores on this scale indicate stronger beliefs in paranormal phenomena ^31^. The internal consistency coefficient for this scale was 0.87, indicating high reliability.

### Stimuli

The visual search task employed a paradigm designed to investigate distinctions between search processes ^32,33^. Commencing each task involved fixing the gaze on a central reference point displayed on the screen. The trial’s target was indicated by briefly superimposing the fixation cross with the target itself, succeeded by the presentation of an array of stimuli incorporating both the target and distractors (Fig. 1A). The task featured a conjunction-style search array, wherein the target shared a common feature property with the distractors, either in terms of shape (‘T’ or ‘L’) or color (red or green) which appeared in one of the 81 locations on the screen. The search arrays varied in size, comprising a total of 6, 12, 24, or 48 stimuli. Participants were tasked with acquiring and maintaining fixation cross on the target until the spacebar was pressed. Their objective was to signal the presence or absence of the target within the array by pressing one of two buttons. Notably, the target was present in half of the trials (Fig. 1A).

### Procedure

The experiment was conducted using Inquisit (version 6.6.1) on a 19-inch LCD monitor. Participants were seated approximately 60 cm from the monitor. Each trial began with the display of a central cross on the screen. Subjects were explicitly instructed to fixate on the dot and press a button to indicate their readiness to proceed. In trials where the target appeared 50% of the time, subjects indicated their decision regarding the target’s presence or absence by manually pressing a button, which then terminated the display. To familiarize themselves with the task and receive performance feedback, each subject completed approximately 24 training trials. Notably, feedback on trial performance was not provided during the testing phase. The task also included an initial fixation cross, which remained on the screen until the participant pressed the spacebar. Following this fixation period, the target for the current trial was briefly presented for 1000 milliseconds. Subsequently, the search array was displayed for 7000 milliseconds. To fulfill the task requirements, each participant completed around 240 trials. They were instructed to respond as quickly as possible without making errors. The entire task lasted for 25 minutes. This study commenced on 01/06/2023 and concluded on 01/04/2024.

### Statistical tests and significance levels

A one-way ANOVA test was conducted to assess the impact of beliefs and set size on search time and accuracy for both TP and TA trials. Search time and accuracy were compared using t-tests for both TP and TA trials. The significance level for all statistical analyses in this study was set at p < 0.05.

## Results

In our investigation into the performance of participants within the conjunction visual search task (Fig. 1A), conducted concurrently with our study, we meticulously analyzed their psychophysical capabilities, considering both the time at which they searched and their accuracy in doing so. This comprehensive study encompassed individuals with paranormal beliefs as well as skeptics (Fig. 1B), allowing for a diverse range of perspectives to be examined. During the task, participants were given specific instructions to identify a designated target amidst a field of distractors following a cue presentation, a trial referred to as target-present trials. Alternatively, they were tasked with signaling the absence of a target if none was present, which constituted target-absent trials (Fig. 1A).

### Believers are fast but careless compared to skeptics in rejecting the presence of the target

In TP trials, no significant difference in search time was observed between believers and skeptics (F_1,3744_ = 4.862, p = .02; Fig. 1C, Fig. 1D). However, in TA trials, believers displayed significantly faster search times when rejecting the presence of the target compared to skeptics (F_1,4150_ = 77.841, p = .001; Fig. 1C, Fig. 1D). Search time showed a significant dependence on set size in both TA (F_3,4150_ = 1335.250, p = .001; Fig. 1C, Fig. 1D) and TP trials (F_3,3744_ = 367.688, p = .001; Fig. 1C, Fig. 1D), with both believer and skeptic groups exhibiting increased search times with higher set sizes in both trial types. Additionally, a tendency toward a significant interaction effect between participants’ beliefs and set size was observed in TP trials (F_3,3744_ = 2.361, p = .06), while a significant interaction effect was observed in TA trials (F_3,4150_ = 8.008, p = .001; Fig. 1C, Fig. 1D). Notably, believers demonstrated faster search times as set size increased compared to skeptics in TA trials.

The accuracy significantly surpassed among skeptics compared to believers across all set sizes in TA trials (F_1,4394_ = 21.216, p = .001; Fig. 1C) and TP trials (F_1,4447_ = 7.445, p = .006; Fig. 1C), indicating that believers are less accurate than skeptics in both finding the target and rejecting its presence. Furthermore, the error rate demonstrated significant dependency on set size in TP trials (F_3,4447_ = 59.803, p = .001; Fig. 1C), whereas no significant dependency on set size was observed in TA trials (F_3,4394_ = 1.652, p = .17; Fig. 1C). Essentially, both groups displayed a decrease in error rate with higher set size in TP trials. Additionally, no significant interaction effect was noted between participants’ beliefs (believers and skeptics) and set size in either TA (F_3,4394_ = 1.992, p = 0.11) or TP trials (F_3,4447_ = 1.589, p = 0.19; Fig. 1C).

In our study, we observed that believers tended to exhibit a propensity for swift responses, indicative of rapid search times. However, this accelerated pace was accompanied by a tendency towards carelessness, particularly evident when discerning the absence of the target stimulus. In contrast, skeptics, while displaying slower search times, demonstrated a commendable level of precision in both rejecting the presence of the target and effectively identifying its absence. This juxtaposition suggests a nuanced interplay between response speed and accuracy, with believers prioritizing rapidity but potentially sacrificing thoroughness, while skeptics adopt a more deliberate approach, resulting in heightened precision.

## Discussion

The observed difference in performance between paranormal believers and skeptics in a conjunction visual search task raises intriguing questions regarding the underlying brain mechanisms contributing to these contrasting behaviors. The finding that paranormal believers tend to search quickly but also exhibit carelessness in rejecting the presence of the target suggests potential differences in cognitive processing compared to skeptics.

One possible explanation for the faster search times among paranormal believers could be attributed to heightened sensitivity or attention toward stimuli associated with their beliefs. Previous research has suggested that individuals with strong beliefs in paranormal phenomena may possess heightened perceptual sensitivity or an increased propensity to detect patterns in ambiguous stimuli ^34,35^. This heightened sensitivity could lead to quicker responses to visual stimuli, including those encountered in a conjunction visual search task. However, the observed carelessness among paranormal believers in rejecting the presence of the target may reflect a cognitive bias or predisposition towards accepting ambiguous or uncertain information. Research has indicated that individuals with paranormal beliefs may exhibit a confirmation bias, wherein they are more likely to accept information that confirms their pre-existing beliefs while disregarding or dismissing contradictory evidence ^36^. This cognitive bias could manifest as a tendency to overlook or misjudge evidence contrary to their beliefs, resulting in a higher rate of false positives or errors in rejecting the presence of the target. In contrast, skeptics’ slower search times but higher accuracy may suggest a more cautious and analytical approach to information processing. Skeptics are less likely to rely on intuition or preconceived notions and may engage in more deliberate and systematic processing of visual stimuli. This analytical approach could lead to longer search times but greater accuracy in distinguishing between target-present and target-absent trials.

Previous studies have provided insights into the brain mechanisms underlying belief processing and decision-making. Functional neuroimaging studies have shown that belief-related processing involves the activation of regions such as the prefrontal cortex, anterior cingulate cortex, and insula, which are implicated in cognitive control, error monitoring, and decision-making ^37,38^. Differences in the activation patterns or connectivity within these brain regions between paranormal believers and skeptics could potentially contribute to the observed differences in visual search performance.

Lastly, the study has encountered several limitations. Firstly, the sample size in the present study was small. Secondly, conducting brain marker investigations via neuroimaging techniques during the conjunction visual search task for both believers and skeptics groups was not feasible. Future studies may yield valuable insights by addressing these constraints.

In conclusion, the contrasting performance of paranormal believers and skeptics in our visual search study highlights potential differences in cognitive processing and decision-making strategies. Future research employing neuroimaging techniques and cognitive neuroscience approaches could provide further insights into the brain mechanisms underlying these behavioral differences.

## Ethical consideration

Our research strictly followed the ethical guidelines outlined in the Declaration of Helsinki. Before participating in the study, all participants provided written informed consent.

## Author contributions

**FA**: Writing – review & editing, Resources, Investigation, Formal analysis. **SA**: Resources, Investigation, Formal analysis. **AN**: Supervision, Formal analysis, Conceptualization, Methodology, Investigation, Project administration, Resources, Writing – original draft, Writing – review & editing.

## Data Availability

All relevant data are available from the corresponding author upon reasonable request.

## Declarations

## Ethics approval and consent to participate

This study was supported by the ethics committee at the Institute for Research in Fundamental Sciences (IPM, School of Cognitive Science), and it was conducted following the ethical standards of the Declaration of Helsinki. All participants gave their informed consent before their inclusion in the study.

## References

1 Tobacyk, J. & Milford, G. Belief in paranormal phenomena: Assessment instrument development and implications for personality functioning. Journal of personality and social psychology 44, 1029 (1983).

2 Wilson, J. A. Reducing pseudoscientific and paranormal beliefs in university students through a course in science and critical thinking. Science & Education 27, 183–210 (2018).

3 Narmashiri, A., Hatami, J. & Khosrowabadi, R. The role of dual mechanism control in paranormal beliefs: Evidence from behavioral and electrical stimulation studies. Cogent Psychology 11, 2316415 (2024).

4 Narmashiri, A., Akbari, F., Sohrabi, A. & Hatami, J. Conspiracy beliefs are associated with a reduction in frontal beta power and biases in categorizing ambiguous stimuli. Heliyon 9 (2023).

5 Narmashiri, A., Sohrabi, A. & Hatami, J. Paranormal beliefs are driving the bias seen in the classification of ambiguous stimuli in perceptual decision-making paradigm. (2023).

6 Narmashiri, A., Hatami, J., Khosrowabadi, R. & Sohrabi, A. Paranormal believers show reduced resting EEG beta band oscillations and inhibitory control than skeptics. Scientific Reports 13, 3258 (2023).

7 Narmashiri, A., Hatami, J., Khosrowabadi, R. & Sohrabi, A. Resting-State Electroencephalogram (EEG) coherence over frontal regions in paranormal beliefs. Basic and Clinical Neuroscience 13, 573 (2022).

8 Narmashiri, A. & Hatami, J. Effect of transcranial direct current stimulation on improving cognitive control in paranormal believers. Journal of Psychological Science 20, 317–326 (2021).

9 Narmashiri, A., Sohrabi, A. & Hatami, J. Brainwave pattern in paranormal beliefs: An EEG-based study in Severe and Mild groups. Neuropsychology 5, 89–98 (2020).

10 Narmashiri, A., Sohrabi, A., Hatami, J., Amirfakhraei, A. & Haghighat, S. Investigating the role of brain lateralization and gender in paranormal beliefs. Basic and Clinical Neuroscience 10, 589 (2019).

11 Narmashiri, A., Sohrabi, A. & Hatami, J. Perceptual processing in paranormal beliefs: A study of reaction time and bias. Social Cognition 6, 113–124 (2018).

12 Narmashir, A. Perceptual-Cognitive Biases in Relation to Paranormal Beliefs: A comparative study in Brain lateralization groups. Neuropsychology 2, 79–92 (2017).

13 Irwin, H. J. The psychology of paranormal belief: A researcher’s handbook. (Univ of Hertfordshire Press, 2009).

14 Wain, O. & Spinella, M. Executive functions in morality, religion, and paranormal beliefs. International Journal of Neuroscience 117, 135-146 (2007).

15 Narmashiri, A., Hatami, J., Khosrowabadi, R. & Sohrabi, A. The role of cognitive control in paranormal beliefs: a study based on performance in go/no-go task. Basic and Clinical Neuroscience 14, 411 (2023).

16 Hergovich, A. & Arendasy, M. Critical thinking ability and belief in the paranormal. Personality and individual Differences 38, 1805–1812 (2005).

17 Musch, J. & Ehrenberg, K. Probability misjudgment, cognitive ability, and belief in the paranormal. British Journal of Psychology 93, 169-177 (2002).

18 Richards, A., Hellgren, M. G. & French, C. C. Inattentional blindness, absorption, working memory capacity, and paranormal belief. Psychology of Consciousness: Theory, Research, and Practice 1, 60 (2014).

19 Dagnall, N., Parker, A. & Munley, G. Paranormal belief and reasoning. Personality and Individual Differences 43, 1406–1415 (2007).

20 Sarbin, T. R. The role of imagination in narrative construction. Narrative analysis: Studying the development of individuals in society, 5–20 (2004).

21 Kaufman, Y., Anaki, D., Binns, M. & Freedman, M. Cognitive decline in Alzheimer disease: Impact of spirituality, religiosity, and QOL. Neurology 68, 1509–1514 (2007).

22 Kraal, A. Z., Sharifian, N., Zaheed, A. B., Sol, K. & Zahodne, L. B. Dimensions of religious involvement represent positive pathways in cognitive aging. Research on aging 41, 868–890 (2019).

23 Georgiou, N., Delfabbro, P. & Balzan, R. Conspiracy beliefs in the general population: The importance of psychopathology, cognitive style and educational attainment. Personality and Individual Differences 151, 109521 (2019).

24 Ballová Mikušková, E. Conspiracy beliefs of future teachers. Current Psychology 37, 692–701 (2018).

25 Van der Wal, R. C., Sutton, R. M., Lange, J. & Braga, J. P. Suspicious binds: Conspiracy thinking and tenuous perceptions of causal connections between co-occurring and spuriously correlated events. European journal of social psychology 48, 970–989 (2018).

26 Van Prooijen, J. W., Douglas, K. M. & De Inocencio, C. Connecting the dots: Illusory pattern perception predicts belief in conspiracies and the supernatural. European journal of social psychology 48, 320–335 (2018).

27 Su, Y., Lee, D. K. L., Xiao, X., Li, W. & Shu, W. Who endorses conspiracy theories? A moderated mediation model of Chinese and international social media use, media skepticism, need for cognition, and COVID-19 conspiracy theory endorsement in China. Computers in Human Behavior 120, 106760 (2021).

28 Denovan, A., Dagnall, N., Drinkwater, K., Parker, A. & Neave, N. Conspiracist beliefs, intuitive thinking, and schizotypal facets: A further evaluation. Applied Cognitive Psychology 34, 1394–1405 (2020).

29 Georgiou, N., Delfabbro, P. & Balzan, R. Conspiracy-beliefs and receptivity to disconfirmatory information: A study using the BADE task. SAGE Open 11, 21582440211006131 (2021).

30 Irwin, H. J. Belief in the paranormal: A review of the empirical literature. Journal of the american society for Psychical research 87, 1–39 (1993).

31 Blackmore, S. & Moore, R. Seeing things: Visual recognition and belief in the paranormal. European Journal of Parapsychology 10, 91-103 (1994).

32 Motter, B. C. & Simoni, D. A. Changes in the functional visual field during search with and without eye movements. Vision research 48, 2382–2393 (2008).

33 Akbari, F., Asivandzadehchaharmahali, S. & Narmashiri, A. Target location and age-related dynamics affect conjunction visual search dynamics. bioRxiv, 2024.2003. 2028.587192 (2024).

34 Lawrence, E. J., Shaw, P., Baker, D., Baron-Cohen, S. & David, A. S. Measuring empathy: reliability and validity of the Empathy Quotient. Psychological medicine 34, 911–920 (2004).

35 Van Elk, M. Paranormal believers are more prone to illusory agency detection than skeptics. Consciousness and cognition 22, 1041-1046 (2013).

36 Lindeman, M. & Aarnio, K. Superstitious, magical, and paranormal beliefs: An integrative model. Journal of research in Personality 41, 731-744 (2007).

37 Van Elk, M., Rutjens, B. T., van der Pligt, J. & Van Harreveld, F. Priming of supernatural agent concepts and agency detection. Religion, Brain & Behavior 6, 4–33 (2016).

38 Kapogiannis, D., Barbey, A. K., Su, M., Krueger, F. & Grafman, J. Neuroanatomical variability of religiosity. Plos one 4, e7180 (2009).

